# Dissecting neural computations of the human auditory pathway using deep neural networks for speech

**DOI:** 10.1101/2022.03.14.484195

**Authors:** Yuanning Li, Gopala K. Anumanchipalli, Abdelrahman Mohamed, Junfeng Lu, Jinsong Wu, Edward F. Chang

## Abstract

The human auditory system extracts rich linguistic abstractions from the speech signal. Traditional approaches to understand this complex process have used classical linear feature encoding models, with limited success. Artificial neural networks have recently achieved remarkable speech recognition performance and offer potential alternative computational models of speech processing. We used the speech representations learned by state-of-the-art deep neural network (DNN) models to investigate neural coding across the ascending auditory pathway from the peripheral auditory nerve to auditory speech cortex. We found that representations in hierarchical layers of the DNN correlated well to neural activity throughout the ascending auditory system. Unsupervised speech models achieve the optimal neural correlations among all models evaluated. Deeper DNN layers with context-dependent computations were essential for populations of high order auditory cortex encoding, and the computations were aligned to phonemic and syllabic context structures in speech. Accordingly, DNN models trained on a specific language (English or Mandarin) predicted cortical responses in native speakers of each language. These results reveal convergence between representations learned in DNN models and the biological auditory pathway and provide new approaches to modeling neural coding in the auditory cortex.

## Introduction

Speech perception involves computations that transform acoustic signals into linguistic representations. When listening to speech, the acoustic signal activates nearly the entire auditory pathway in the human brain: from the auditory nerve and subcortical structures to primary and nonprimary auditory cortices. Natural speech perception is a challenging task with considerable variability in the acoustic cues for the same linguistic perceptual units such as phonemes, syllables, and words. This is due to varying contextual factors such as different speakers, emotions, prosody, coarticulation, speech rate, and lexical context.^1–4^ Despite these challenges, the human auditory system is sensitive to this variability yet robustly extracts invariant representations of the phonetic and lexical information to support speech comprehension.^3,5-8^ A central goal of speech auditory neuroscience, as well as cognitive neuroscience in general, is to understand the computations performed in specific neural circuits and the representations generated by such computations.^9^

Classical cognitive models of speech perception, such as the COHORT model,^10^ the TRACE model^11^ and their variants, account for many psychological aspects of speech perception, but do not explain neural coding or perform well for natural speech recognition. On the other hand, classical neural encoding models ^12–15^ explain neural coding during speech perception, but cannot be directly adapted to a unified computational framework of speech perception. Only recently have modern artificial intelligence (AI) models using deep neural networks (DNN) finally approached human-level performance in automatic speech recognition.^16–19^ However, these data-driven models are trained as end-to-end “black boxes” and it is not clear how to interpret the computations they implement and the representations they generate. Here, we are interested in understanding how the computations and representations of these DNN models relate to those found in the human auditory system. Connecting AI models to neural coding of human sensory systems has important implications on their interpretability and offers new data-driven approaches toward computational models of sensory perception.

Task-oriented pre-trained DNN models have shown promise as computational models for sensory neuroscience. In particular, using the internal representation features learned from supervised learning tasks, such as image recognition or sound classification, encoding models predict evoked neural population responses in visual and auditory cortices with high accuracy.^20–24^ Two of the key ingredients in neural network models are the model architecture and the training objective. Model architecture determines the type of computations performed on input signals, while the training objective prioritizes specific types of representations that the model is learning through optimization. The neural coding in the ventral visual cortex is largely driven by spatial statistics in the retinotopic space,^25^ therefore convolutional neural networks built upon a hierarchy of spatial convolutions have had great success in modeling neural representations in the ventral visual pathway.^20,23,24^

Unlike core object recognition in vision modeling which uses static images, speech is inherently dynamic and can be better modeled with sequence-to-sequence (seq2seq) learning models rather than spatial pattern recognition. Furthermore, supervised model training, which often requires an enormous amount of labeled data, is not plausible as a generic learning strategy of the human auditory system. Human infants can learn phonetic and linguistic categories from the statistical distribution of speech sounds in native languages without explicit word learning.^26–28^ Recent works have suggested that unsupervised models, which do not require labeled data, can also be used as models of vision and high-level language processing in brain.^29–31^ Therefore, (unsupervised) speech models which learn the transient (local) features of speech as well as the statistics of longer sequences and context, may yield more optimal models for speech perception.^32^

In this study, we directly assess the representational and computational similarities between the state-of-the-art speech neural network models and the human auditory pathway in order to shed light upon the underlying shared computations in the two systems. On the biological side, neural responses to natural speech in different parts of the ascending auditory pathway are extracted from biophysical models determined from neural recordings and direct intracranial electrophysiological recordings from the human auditory cortex; on the DNN side, different levels of speech embeddings are extracted from pretrained speech models. Using neural encoding framework,^12,33^ we systematically evaluate the similarity between the auditory pathway and DNN models with different computational architectures, such as convolution, recurrent and self-attention, and with different training strategies, including both supervised and unsupervised objectives. Furthermore, by inspecting the context-dependent computations in the DNNs, we provide interpretable insights into the underlying sequential and contextual computations and representations that drive the predictions in the neural encoding models. Unlike previous modeling efforts which mainly focused on one single language, mainly English, we use a cross-linguistic paradigm and test if our DNN based models can reveal language-invariant and language-specific aspects during speech perception.

In particular, we demonstrate the following findings: 1) the hierarchy in the DNNs trained to learn speech representations correlate to the ascending auditory pathway; 2) unsupervised models without explicit linguistic knowledge are able to learn similar feature representations as the human auditory pathway; 3) deeper layers in speech DNN correlate to the context-dependent speech-responsive populations in non-primary auditory cortex, and the correlation can be explained by the specific computations aligned to important linguistically-relevant temporal structures in speech, such as phonemic and syllabic contexts; 4) the DNN-based model is able to reveal language-specific properties in cross-language speech perception, which is not easily captured by traditional linear encoding model. Taken together, we provide new data-driven approaches to modeling and evaluating neural coding in the auditory cortex.

## Results

### Overview

Our overall goal is to understand computations and representations that occur and emerge throughout the auditory system during speech perception. To model the early portion of the pathway, we used a biophysical model simulation from the auditory periphery and midbrain,^34–36^ which have been highly successful as neural encoding models at the cellular level. The biophysical model yielded 50 distinct neurons in the auditory nerve (AN) and 100 distinct neurons in the inferior colliculus (IC). For the later portion of the pathway, we used intracranial cortical recordings from the human auditory cortex, including both primary and non-primary cortices.^37^ High-density grid electrodes were placed on the auditory cortex in nine participants (Supplement Fig. 1) and local field potentials were recorded as these participants listened to an English speech corpus. From the total of 553 electrode sites along the auditory cortex, 81 electrodes were located in the primary auditory cortex (Heschl’s gyrus; HG) and 472 were located in the non-primary auditory cortex (superior temporal gyrus; STG). The amplitude of the local field potential in the high-gamma band (70-150 Hz) was computed and used as a measure of the local neuronal activity.^38^ Neural responses across the early and later auditory systems were assessed using a set of 599 English sentences from the TIMIT corpus.^39^

We used four deep neural networks as computational models to extract speech representations from speech stimuli. A critical factor that differentiates these models is their training objectives. In particular, we employed two unsupervised models and two supervised models: 1) the HuBERT model is a transformer-based self-supervised model trained to perform prediction of masked portions in speech;^19^ 2) the Wav2Vec 2.0 unsupervised model (W2V unsup) is a transformer-based self-supervised model trained to perform a contrastive learning task that distinguishes spans of the speech utterance from distractors;^17^ 3) the Wav2Vec 2.0 supervised model (W2V ASR) is a transformer-based supervised model based upon fine-tuning the unsupervised model using automatic speech recognition objective;^17^ 4) the Deep Speech 2 model (DS2) is an LSTM-based supervised model trained with automatic speech recognition objective.^16^ These models share a similar hierarchical framework composed of a multi-layer convolutional feature encoder and a multi-layer sequential context representation encoder. The convolutional feature encoder takes raw speech audio (raw waveform or spectrogram) as input and extracts latent speech representations using 1D and 2D convolutions. These latent representations are hypothesized to reflect temporally constrained lower-level acoustic features. The sequential context representation encoder, which consists of multiple Transformer encoder layers or multiple recurrent layers (long short-term memory layers), takes in the output from the convolutional feature encoder and extracts contextual information from the sequence, which is hypothesized to reflect higher-level context dependent phonetic information. We pre-trained the speech learning models on Librispeech, a standard corpus of 960h continuous naturalistic English speech.^40^ (Table 1)

**Table 1.** Summary of network training objectives, architectures, and dimensionality.

| Models | Unsupervised objective | Supervised objective | Architecture | # of dimensions in each layer |
| --- | --- | --- | --- | --- |
| HuBERT (Hsu et al., 2021) | Masked prediction | N/A | 7 CNN layers + 12 Transformer encoder layers | 512 (CNN)<br>768 (Transformer) |
| Wav2Vec 2 (unsupervised) (Baevski et al., 2020) | Contrastive learning | N/A | 7 CNN layers + 12 Transformer encoder layers | 512 (CNN)<br>768 (Transformer) |
| Wav2Vec 2 (Supervised) (Baevski et al., 2020) | Contrastive learning | Automatic speech recognition | 7 CNN layers + 12 Transformer encoder layers | 512 (CNN)<br>768 (Transformer) |
| Deep Speech 2 (Amodei et al., 2016) | N/A | Automatic speech recognition | 3 CNN layers + 5 LSTM layers | 1312 (CNN)<br>1024 (LSTM) |

The speech responses from both the auditory pathway and DNNs were aligned in time, and linear encoding models were trained using different representation layers in the DNNs to predict neural responses in the auditory pathway (Fig 1). The performance of these neural encoding models (prediction R^2^) quantifies the similarity between the learned speech representation in the DNNs and the underlying neural representation in the auditory pathway. In this way, we test the hypothesis that the speech DNN models converge to a similar representation hierarchy as the ascending auditory pathway.

**Figure 1.**
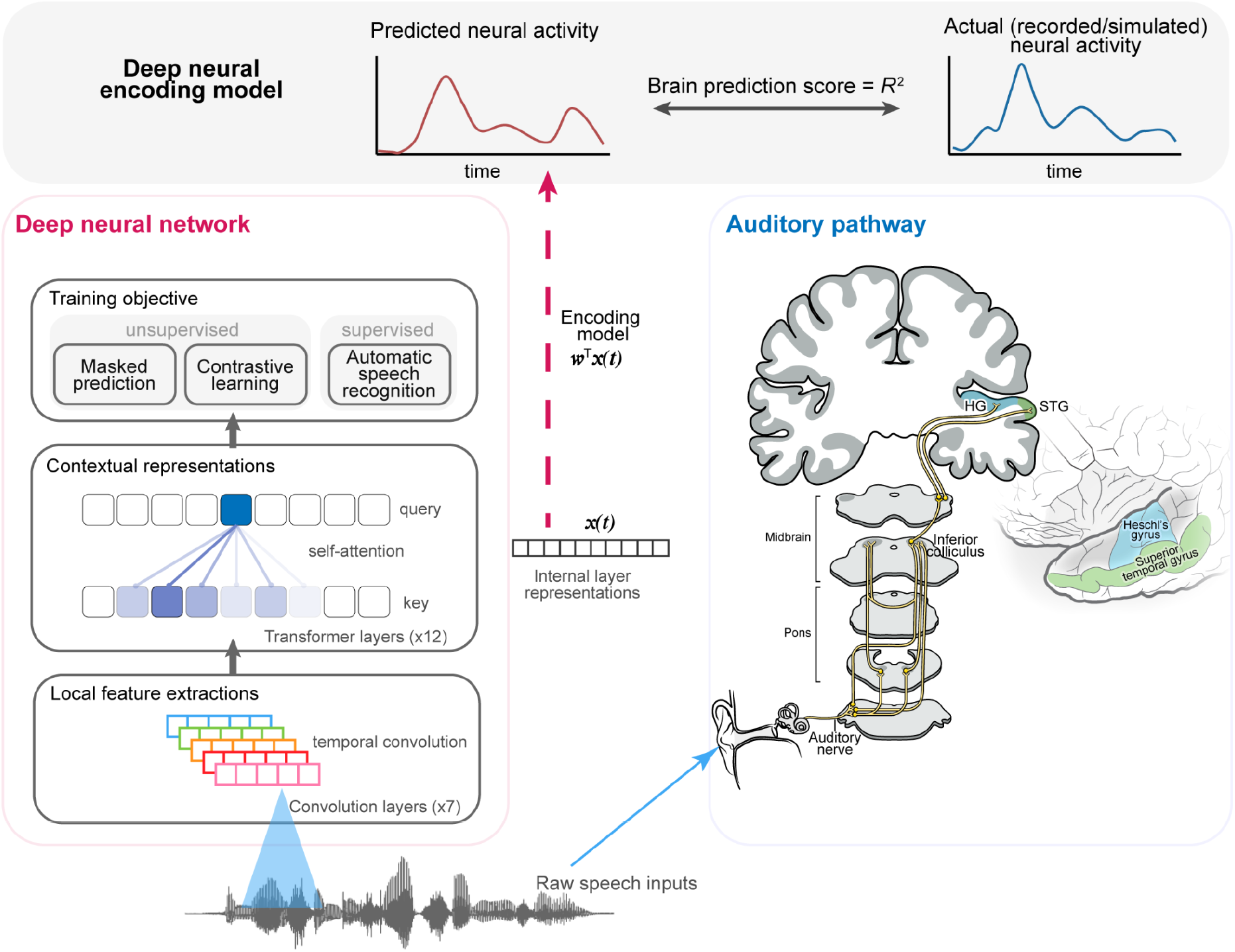
The overall framework for comparing representations in deep neural networks and the auditory pathway. The architecture of a family of deep neural network models, the Wav2Vec2.0/HuBERT, is illustrated on the left. The auditory pathway is illustrated on the right, with highlighted areas indicating the locations of the recorded/simulated electrophysiology signals. The same natural speech stimuli were presented to both the human subjects and the DNN models, and the internal activations of each DNN layer were extracted and aligned to the corresponding neural activity from each recording site of the auditory pathway. A ridge regression model was fitted to predict the neural activity from time-windowed DNN representations, and the correlation *R*^2^ between the predicted and actual neural activity was used as a metric of prediction accuracy (brain prediction score; BPS).

To account for the heterogeneous signal-to-noise ratio in different parts of the auditory pathway, we first set up benchmark baselines for each individual electrode and neuron in our recordings. Specifically, for each recording site, we trained two baseline models: 1) a linear temporal receptive model using spectrogram features;^12^ 2) a linear temporal receptive field model using a heuristic set of acoustic-phonetic features that include spectrogram, speech envelope, and temporal landmarks, pitch, and phonetic features.^37^ (Supplement Fig 2) The performance of the neural encoding models using DNN features was normalized to these baselines in each individual recording site in order to make evaluations and comparisons across recording sites and areas. This normalized prediction R^2^ is termed as the brain prediction score (BPS), and used as the major metric for prediction accuracy for each recording site.

### The hierarchy of layers in DNNs correlate with the ascending auditory pathway

We test whether DNNs that are trained to learn speech representation converge on the same standard auditory serial feedforward hierarchy of AN-IC-HG-STG. To do this, we compared the DNN hierarchy and the ascending auditory pathway from three different perspectives: 1) are the feature representations learnt by DNNs more strongly correlated with the neural coding than linguistically-derived acoustic-phonetic feature sets? 2) Does the hierarchy of layers in DNNs mirror a similar hierarchy in the ascending auditory pathway? 3) Do DNN encoding models reflect different temporal integration profiles along the auditory pathway?

First we took a representative state-of-the-art self-supervised DNN, the HuBERT model.^19^ For each single-layer representation model in HuBERT, we computed the averaged brain prediction score (normalized prediction R^2^) across all recording sites within each anatomical area (Fig 2a). Compared to the linear model with heuristic acoustic-phonetic features, the DNN encoding model explained 39.9% more variance in AN at Transformer layer 1 (t(50) = 13.97, p = 2.5e-44, two-sided), 76.3% more variance in IC at Transformer layer 1 (t(100) = 13.75, p = 5e-43, two-sided), 3.4% more variance in HG at Transformer layer 1 (t(53) = 1.20, p = 0.23, two-sided), and 23.0% more variance in STG at Transformer layer 10 (t(144) = 16.1, p = 5e-58) (Fig 2a). Moreover, out of all layers in the same unsupervised DNN model, AN and IC responses were best predicted by the CNN layers as well as the first 4 Transformer layers in the hierarchy (Fig. 2a). The speech responsive STG population was best predicted by the later part of the DNN model, and peaked at the 10th layer out of all 12 Transformer layers (Fig. 2a). HG responses were predicted by all Transformer layers equally well. However, none of these speech DNN layers out-performed the baseline acoustic model for HG (Fig. 2a).

**Figure 2.**
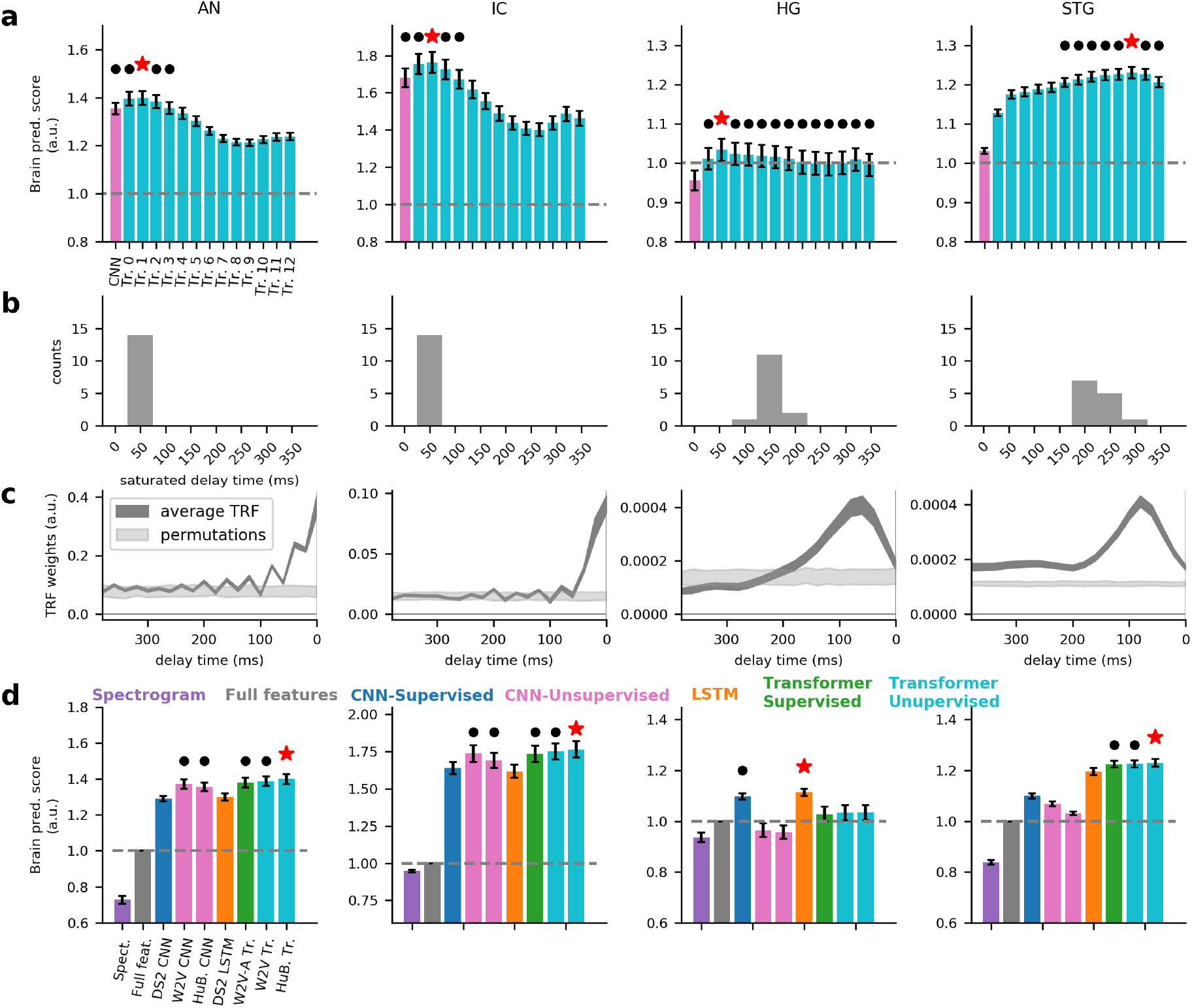
The hierarchy of layers in DNNs correlate to the AN-Midbrain-STG ascending auditory pathway. **a)** The averaged normalized brain prediction score of the best-performing neural encoding model based on each single layer in the HuBERT model (maximum over delay window length). Magenta bars indicate CNN output layers, cyan bars indicate Transformer layers. Red star (*) indicates the best model for each area, black dot (.) indicates other models that are not statistically different from the best model (p > 0.05, two-sided paired t-test), from left to right: AN, IC, HG and STG, same for each row in **b-d. b)** The histogram of the optimal delay window lengths corresponding to models in **a. c)** The averaged temporal receptive field (absolute beta weights of the spectrotemporal encoding model) in speech-responsive units/electrodes of each area. (mean ± s.e.m., light shaded areas indicate random permuted distributions). **d)** The averaged normalized brain prediction score of the best-performing neural encoding model types (maximum over single layers and delay window lengths) for different areas of the pathway. Color key indicates different layer type. Red star (*) indicates the best model for each area, black dot (.) indicates other models that are not statistically different from the best model (p > 0.05, two-sided paired t-test). Dashed horizontal line indicates baseline model using full acoustic-phonetic features.

Next, we tested the hypothesis that the auditory hierarchy is characterized by increasingly long temporal integration windows. The Transformer-encoding models demonstrated distinct temporal integration profiles for different areas: AN and IC encoding models all showed dominant delay time window length of 50 ms. HG delay time window lengths were around 100-200 ms, while STG delay time window lengths were larger than 200 ms (Fig 2b). Using the baseline spectrogram model, we found the temporal receptive fields estimated for each area showed a hierarchy of progressive temporal integration of acoustic inputs: temporal responses in peripheral areas AN and IC were mostly transient within 100ms, while neural responses in the cortex showed integration time windows longer than 100ms. More specifically, HG on average had a consistent temporal integration window of 200 ms, while STG showed more diversified profiles, some with transient integration windows less than 200ms and some with significant sustained temporal integration longer than 400ms (Fig 2c).

Finally, we generalized the evaluations to a set of different DNN models in Table 1. We found that for all areas, all DNN encoding models outperformed the baseline linear models. On average, compared to the linear model with heuristic acoustic-phonetic features, DNN encoding models explained 29.3%-40.0% more variance in AN, 61.7%-76.3% more variance in IC, −3.5%-11.4% more variance in HG, and 3.1%-23.0% more variance in STG (Fig 2d). In particular, the Transformer layers in the unsupervised HuBERT model achieved highest average performance in all areas except for HG. Moreover, we found that neural responses to speech in the auditory periphery (AN & IC) and primary auditory cortex (HG) were also largely characterized by locally resolved filters such as CNN representations, which had a fixed finite receptive field in time (p > 0.05 compared to HuBERT, two-sided t-test, Fig. 2d). On the other hand, speech responses in the non-primary auditory cortex (STG) were better predicted using context-dependent feature representations layers (Transformer layers) of the DNNs (Fig. 2d), which dynamically track the context information in the sequential speech input.

To sum up from the above three perspectives, the early to later layers in the deep neural networks trained to learn speech representations correlate to the successive processing in the ascending auditory pathway. HG representation is not modeled well by speech DNNs (p > 0.1 in all layers compared to baseline, Fig 1a) despite the latencies and temporal integration window for TRFs would suggest a serial processing pathway.

### Sustained speech responses in STG are explained by deeper contextual layers in DNN

Previous studies have identified neural populations in STG that show distinct speech responsive profiles, including onset and sustained responses.^37,41^ Here, we evaluated whether these functionally distinct speech-responsive populations corresponded to different layers of contextual dependent representations in the same DNN model.

Among the 144 speech-responsive electrodes in STG, we found 2 clusters that showed distinct transient and sustained response profiles, based on averaged high-gamma responses across sentences (Fig 3a, 3b, and Fig S2). We then looked into the best STG-prediction model, the HuBERT model, and compared the brain prediction scores of different layers with regard to these functional clusters. As shown in Fig. 3c, for the more sustained clusters (Clusters 1), the best prediction model came from the deep layers of the Transformer-encoder in the DNN (Clusters 1: peak BPS = 1.26 at Transformer layer 10), which were also significantly higher than the early layers in the DNN (p < 0.05, two-sided paired t-test, df=83, no statistical difference between layers 6-12). For the more transient cluster (Cluster 2), the best prediction model was from the Transformer layer 5 in the DNN (peak BPS = 1.20 at layer 5). However, the peak prediction layer did not significantly outperform any other Transformer layers in the network, only except the very first one (p > 0.05 for all two-sided paired t-test, df=61 for cluster 2). Both clusters 1 and 2 showed a similar optimal delayed-time window length of around 200-250ms (Fig 3d). As a result, the sustained speech-responsive neural activity prevalent in STG can be predicted from the deeper contextual representation layers in DNN, while the more transient speech-responsive neural activity, such as the onset response, can be predicted in both the early and late part of the Transformer hierarchy in DNN. This indicates that these different neural populations do not correspond to a simple serial feedforward process in the DNN hierarchy, where the more transient population is mapped to early layers and the more sustained population is mapped to the later layers. On the other hand, the DNN maintains the transient onset representation throughout the processing hierarchy, and the later layers represent both transient and sustained representations in parallel.

**Figure 3.**
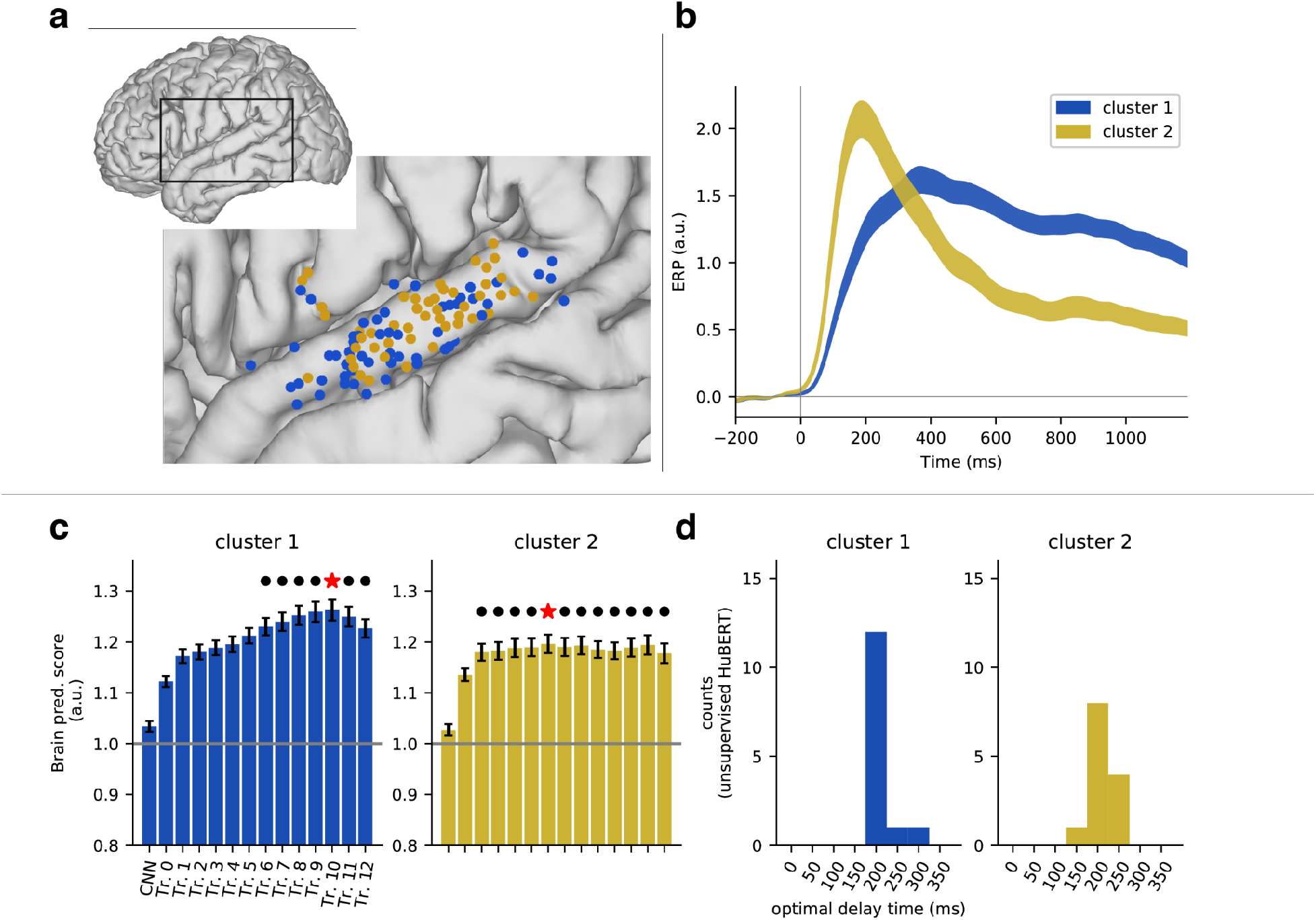
Functional subpopulations in STG correlate to different contextual representation layers in DNN. **a)** Anatomical locations of all speech responsive electrodes, mapped onto a common cortical space. Different colors indicate different functional clusters. **b)** The averaged event-related potential of each functional cluster. All time aligned to sentence onsets, and normalized to resting-state baseline. (mean ± s.e.m) **c)** The averaged normalized brain prediction score of the encoding model based on every single layer in HuBERT for each functional cluster (maximum over delay window length). Red star (*) indicates the layer with highest score, black dot (.) indicates other layers that are not statistically different from the best model (p > 0.05, paired t-test). **d)** The histogram of the optimal delay window lengths corresponding to models in **c**.

### DNN context-dependent processing explains encoding predictions in human cortex

We next examined the computational mechanism underlying contextual representations in the DNN. We asked whether certain types of contextual computation for speech in DNN explain the ability to predict brain responses.

Specifically, we extracted the attention weight matrices in each Transformer layer of the HuBERT model, which quantified the contributions from different parts in the context to the feature representation at each time. Critically, these contextual attention weight matrices were not static filters but rather dynamically changed according to the specific speech sequences. Therefore, it reflected the stimulus-dependent dynamic extraction of contextual information in each speech sequence. Such computations would be critical for resolving contextual effects,^42^ such as allophonic variation,^43^ and extracting invariant representation of lexical information from acoustic signals.

As a result, for each sentence in the speech corpus, we defined templates of attention matrices corresponding to different levels of contextual information representation in speech, including contextual information within the same phoneme, contextual information from previous phoneme(s), contextual information within the same syllable, and contextual information from previous syllable(s) (Fig 4a, b). We then computed the averaged correlation coefficient between the actual attention weight matrices in each DNN layer and the templates across all sentences, which we termed as the attention score for each layer in DNN (Fig 4c). We found a general trend that deeper layers showed an increased amount of contextual attention to linguistic structures (previous phoneme(s) and syllable(s)) (Fig 4c, bar plots). A randomized DNN model with the same architecture but no pre-training on speech data did not show such progressive contextual attention along the hierarchy (Fig 4c, black lines). Therefore, the alignment of attention to contextual structures was not just a direct consequence of the hierarchical architecture of the DNN model that emerges with depth but reflecting computations adapted to extracting speech-specific linguistic-relevant representations through training on natural speech. (Fig 4c).

**Figure 4.**
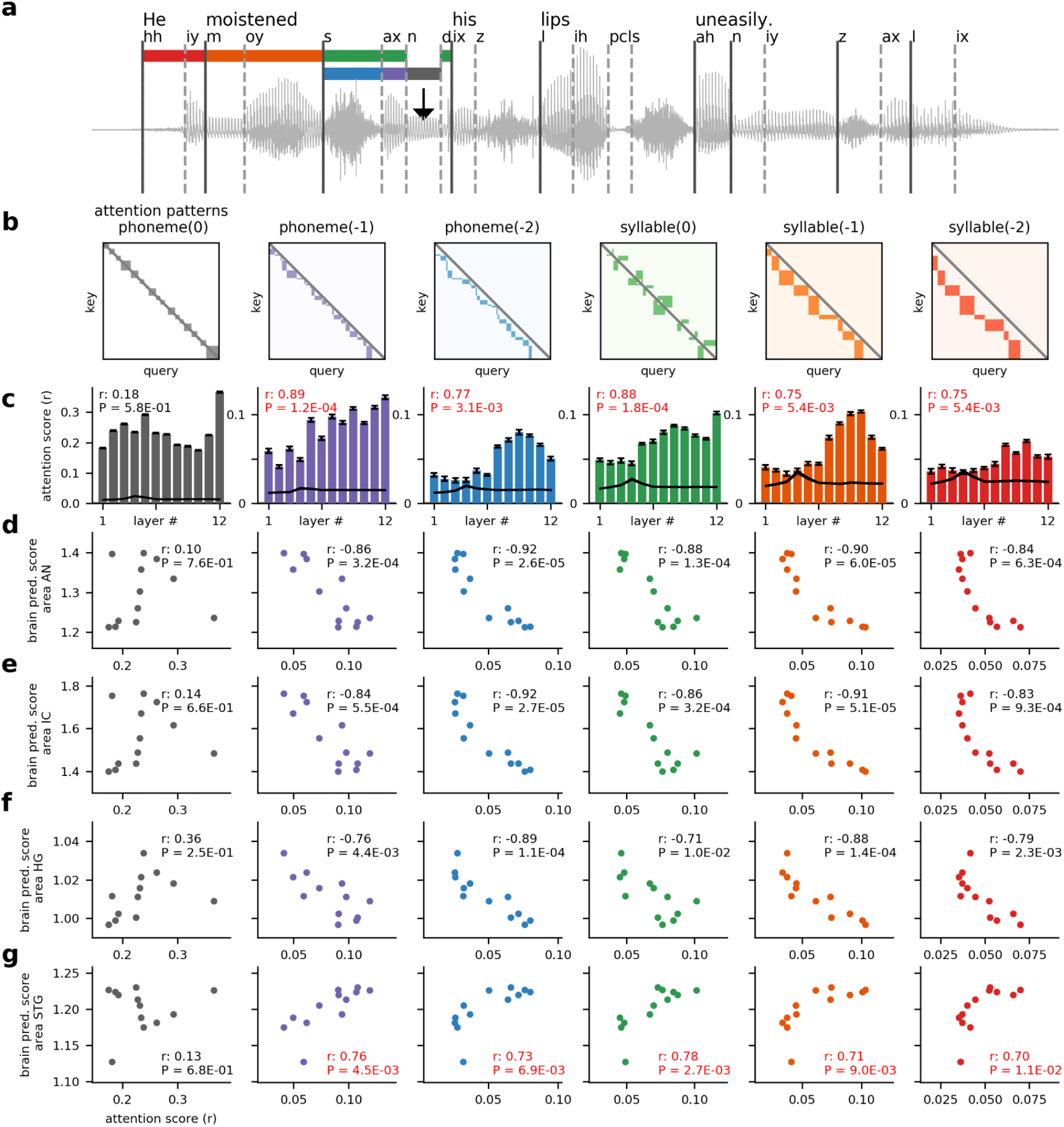
Context-dependent computations explain brain correspondence across layers in DNN. **a)** Sample speech sentence text, waveform, and phonemic annotations are shown. The segmentations of phonemic and syllabic context to the current time frame (black arrow) are marked in different colors. phoneme(0): current phoneme (grey); phoneme(−1): previous phoneme (purple); phoneme(−2): second-to-previous phoneme (blue); syllable(0): current syllable (excluding current phoneme, green); syllable(−1): previous syllable (orange), syllable(−2): second-to-previous syllable (red). **b)** The template attention weight matrices for different contextual structures as shown in a). Query: the target sequence: key: the source sequence. Colored blocks correspond to different contexts. **c)** The averaged attention score (Pearson’s correlation coefficient between attention weights and template) across all English sentences for each Transformer layer corresponding to each type of attention template. r values in the top left of each panel indicate the correlation between attention score and layer index (df = 12). Black line indicating averaged attention score from the same DNN architecture with randomized weights (mean +/- s.e.m.). **d-g)** Scatter plots of attention score vs. brain prediction score across layers, each dot indicates a Transformer layer, each panel corresponds to one type of attention pattern. The r and p values correspond to the AS-BPS correlation across layers (Pearson’s correlation, df = 12). Each row corresponds to a different area (**d** - AN, **e** - IC, **f** - HG, **g** - STG).

We then tested if such trends in contextual computations would predict the brain-prediction performance for different layers in the DNN. Specifically, we correlated the attention score to the brain prediction score for each brain area in different DNN layers. We found that the phonemic and syllabic level attention to the linguistic context in speech was positively correlated to the ability to predict brain activity only in the non-primary auditory cortex (Fig 4g), but not in the auditory periphery or the primary auditory cortex (Fig 4d-f). In other words, for a given Transformer layer in the model, the better the attention weights aligned to linguistic contextual structure, the better the layer’s learned representation would be able to predict speech response in STG. On the other hand, the more contextual information attended, the less the learned representation would be correlated to AN/IC/HG response. Since the latent representation in a Transformer layer is a combination of locally constraint representation and its contextual representations, these results also indicate that neural code in STG is reflective of contextual linguistic information, while the neural code in AN/IC/HG is more reflective of temporally-constrained acoustic representations.

### DNN context-dependent representations capture language-specific information

Next, we tested whether the contextual dependent computation was language-specific and reflected higher-level language processing beyond just the acoustics, such as phonotactic, phonological, or lexical representations. To do this, we took a cross-linguistic approach, comparing English and Mandarin (Fig. 5a). Mandarin shares many of the consonants and vowels of English but differs drastically in how phonetic and prosodic features are combined to give rise to words. In addition to the English-speaking participants, we also analyzed cortical recordings from native Mandarin speakers. Both groups were monolingual and had no comprehension of the foreign language. We adopted the same paradigm and materials as our previous study which mainly focused on cross-linguistic pitch perception^44^. The two groups were instructed to listen to both naturalistic English speech and Mandarin speech in separate recording blocks. In addition to the previous HuBERT model pretrained on English speech, we also pretrained the same HuBERT model with naturalistic Mandarin speech. We then compared the performance of the two models on the two groups when they listened to different languages (Fig. 5a).

**Figure 5.**
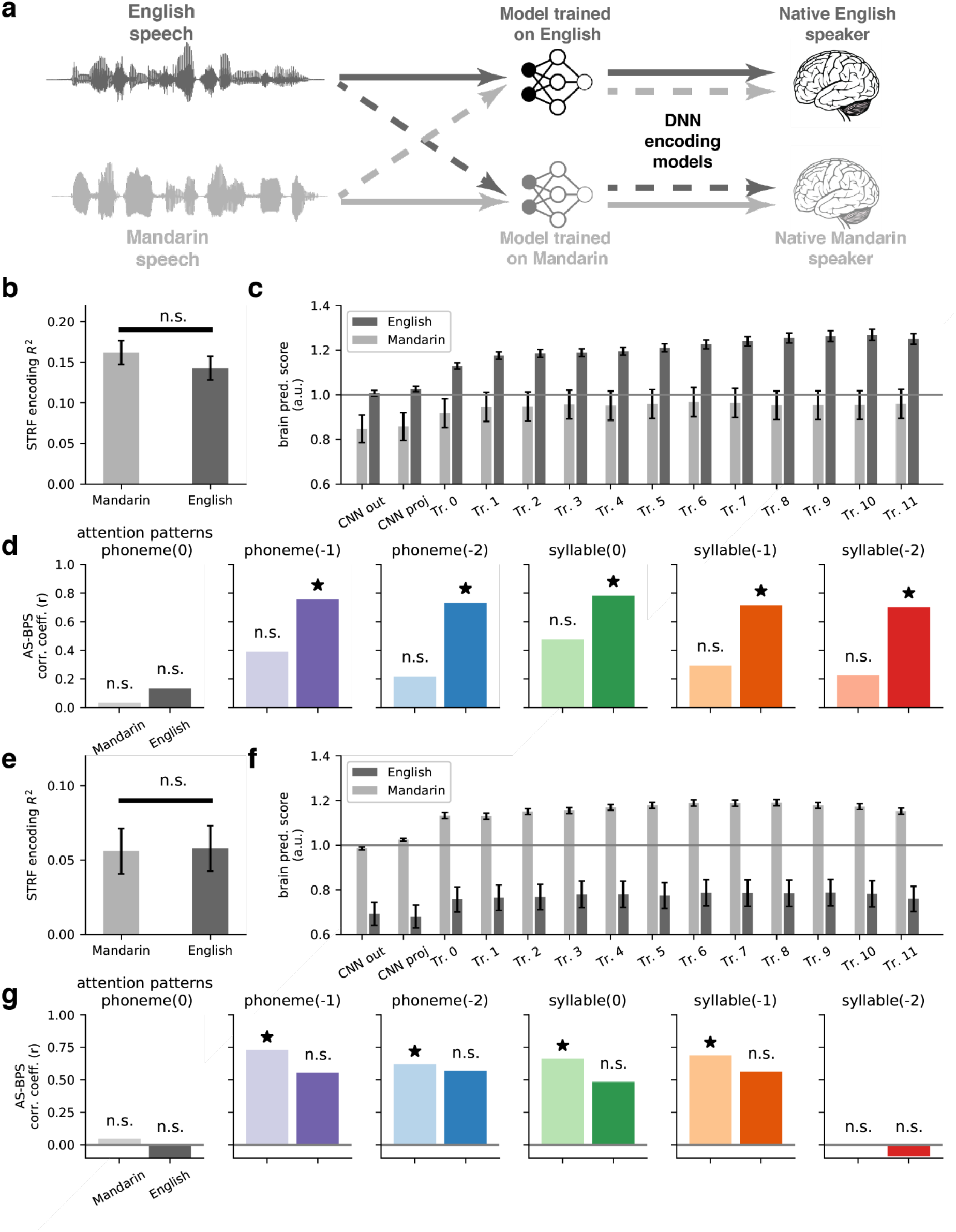
Cross-language encoding comparisons reveal language-specific representation and computations aligned between DNN and STG. **a)** Schematic of the cross-language paradigm. Both English (darker color) and Mandarin (lighter color) speech were fed into models pretrained on English or Mandarin. The extracted representations were used to predict neural responses recorded in STG from native English speakers or native Mandarin speakers when they listened to the corresponding speech. **b)** The averaged prediction R^2^ of linear STRF model in STG electrodes from native English speakers using English or Mandarin speech. Two-sided paired t-test. **c)** The averaged normalized brain prediction score of the encoding model based on every single layer in English-pretrained HuBERT model in native English speakers when listening to English vs Mandarin speech. All comparisons are significant with p < 0.01, paired two-sided t-test. **d)** The AS-BPS correlation across layers in English-pretrained HuBERT model and STG in native English speakers (Pearson’s correlation, df = 12). Each panel corresponds to one type of attention pattern. (See also Fig. 4). **e-g)** Same as **b-d**, but using Mandarin-pretrained HuBERT model and recordings from STG in native Mandarin speakers.

To explicitly test our hypotheses of linguistically relevant context dependent processing in the auditory pathway as shown in the previous section (Fig. 4), we conducted cross-lingual perception and DNN prediction tests. In particular, we hypothesized that the contextual dependent computation in DNN capture language-specific higher-level processing beyond acoustics in STG, and therefore we would expect English-pretrained model shows higher brain prediction performance in STG of native English speakers, and the prediction performance is better aligned to contextual attention to the phonemic and syllabic structures in English than Mandarin. On the contrary, the Mandarin-pretrained model would show higher brain prediction and better correlation to contextual attention for Mandarin speech in native Mandarin speakers.

First, we looked at the results with an English-pretrained model and native English speakers. At the acoustic level, the linear STRF model, which only included spectrogram features, showed similar performance in predicting neural responses in STG when listening to different languages (mean R^2^ = 0.162 and 0.143 for Mandarin and English speech respectively, paired *t*(45) = 1.65, *p* = 0.105, two-sided, Fig 5b). This suggests that the lower-level acoustic representation is largely shared across languages. However, a performance gap was found in the DNN encoding models between languages, where the brain-prediction score for English speech was significantly higher than Mandarin speech (Fig 5c). Moreover, the gap between the two languages monotonically increased in deeper layers of the network. ΔBPS = 0.160 at CNN output layer (p=0.007, paired t-test, two-sided), ΔBPS = 0.211 at the first Transformer encoder layer (p=0.005, paired t-test, two-sided), ΔBPS = 0.314 at the 10th encoder layer (p=2e-5, paired t-test, two-sided) (Fig 5c). This suggests that the representation in the network demonstrates an increasing level of language-specific information. We also evaluated the relationship between the phonemic and syllabic contextual information computation in DNN layers and the corresponding brain-prediction performance for Mandarin speech in STG. As opposed to the previous results (Fig 4g), no significant correlation was found in either phonemic or syllabic level between the attention patterns in DNN layers and brain-prediction scores when listening to Mandarin speech (Fig 5d).

In contrast, we found opposite results with a Mandarin-pretrained model and native Mandarin speakers. At the acoustic level, the linear STRF model also showed similar performance for both Mandarin and English speech (mean R^2^ = 0.056 and 0.058 for Mandarin and English speech respectively, paired *t*(92) = −0.501, *p* = 0.617, two-sided, Fig 5e). The DNN encoding models showed consistently higher performance for neural response to Mandarin speech than English, and the gap also increased in deeper layers. ΔBPS = 0.293 at CNN output layer (p=2e-6, paired t-test, two-sided), and ΔBPS = 0.405 at the 9th Transformer encoder layer (p=5e-8, paired t-test, two-sided) (Fig 5f). Moreover, as opposed to the English-pretrained model and native English speakers combination, we found consistently significant correlations between phonemic or syllabic level attention scores and brain-prediction scores when listening to Mandarin speech, and no significant correlation when listening to English speech, in these native Mandarin speakers (Fig. 5g).

Therefore, our results demonstrated a double-dissociation pattern between pre-trained models and native languages, suggesting that the contextual dependent computations in the DNN model captured higher-level language-specific linguistic information in STG that are learned depending on language experience.

### Acoustic-phonetic feature representation in DNN explains brain prediction performance

The last question we asked is whether the brain-prediction performance of the DNN layers can be accounted for by an acoustic-to-phonetic processing hierarchy. We tested the feature representations of acoustic, phonetic and prosodic information in the DNN layers. Specifically, we applied similar linear feature encoding models to predict the activations of the hidden units in different DNN layers and computed the unique variance explained by each set of features. It is worth pointing out that these features are statically coded and do not vary according to different contexts. Therefore, our analysis here intentionally reflects the static non-contextual part of acoustic/phonetic/prosodic representations in the DNN layers, as addressed in the previous analyses.

Overall, the results indeed demonstrated an acoustic-to-phonetic transformation along the hierarchy (Fig 6A). In the CNN output layer, the spectrogram features uniquely accounted for 20.0% of the total variance, while phonetic features only accounted for 1.70% (paired t(768) = 47.6, p < 1e-10, two-sided). However, after the 3rd Transformer encoder, phonetic features consistently explained more unique variance than the acoustic features in the network. The unique variance explained by static phonetic features peaked at the 11th Transformer encoder layer with unique R^2^ = 3.98% (paired t(768) = 9.12, p < 1e-10, two-sided t-test against acoustic features). On the other hand, temporal landmark features, such as speech envelope and onsets, and prosodic pitch features (absolute and relative pitch) were more uniformly distributed along the hierarchy of the network (Fig 6a).

**Figure 6.**
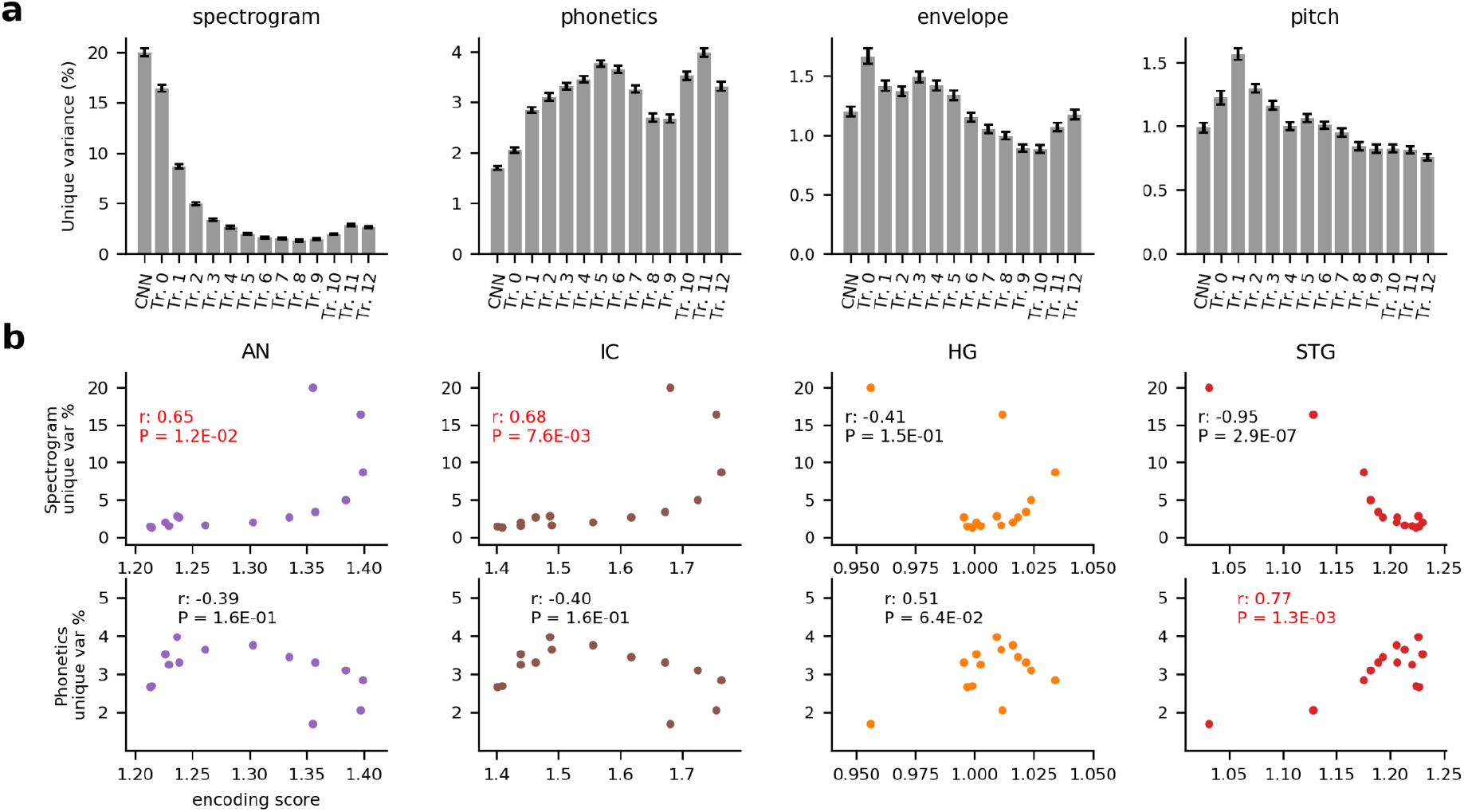
The representations in neural networks demonstrate an acoustic-to-phonetic transformation hierarchy, yet preservation of prosodic cues through DNN layers. **a)** The average unique variance explained by each set of features across units in each DNN layer. **b)** First row: the correlation between brain prediction score and unique variance explained by spectrogram features in each layer. Second row: the correlation between brain prediction score and unique variance explained by phonetic features in each layer. Each panel corresponds to one area. From left to right: AN, IC, HG, STG.

Furthermore, when correlated to the brain-prediction score of individual layers, spectrogram and phonetic feature encoding only showed significant positive correlation in the peripheral areas (AN: Pearson’s r = 0.65, p = 0.012; IC: Pearson’s r = 0.72, p = 0.0039; Fig 6b). Furthermore, the phonetic feature encoding also correlates with brain-prediction score in STG (Pearson’s r = 0.77, p = 0.0013; Fig 6B), but not the other areas (p > 0.05 for all other 3 areas; Fig 6B). Taking these together, a similar acoustic-to-phonetic hierarchy was found and correlated in both the self-supervised DNN model and the ascending AN-IC-STG pathway.

## Discussion

We have demonstrated that computational representations of speech learned in state-of-the-art deep neural networks resemble important aspects of information processing through the human auditory system. The feature representations learned by DNNs significantly outperform theory-driven acoustic-phonetic feature sets in predicting neural responses to natural speech throughout the auditory pathway. We find that the hierarchy of layers in DNNs correlate to the AN-Midbrain-STG ascending auditory pathway. Contextualized representations learned by DNNs correlate to functionally distinct speech-tuned populations in the non-primary auditory cortex. Furthermore, by inspecting the core contextual computation components in DNNs, we demonstrate that DNN models are able to learn critical linguistically-relevant temporal structure, such as phoneme and syllable contexts, from natural speech even with unsupervised training. Such ability to learn language-specific linguistic information also predicts the correlation between the learned representation in DNN and the neural coding in non-primary auditory cortex. The DNN-based neural encoding model is able to reveal language-specific coding in STG during cross-language perception, which is not sensitive

Encoding models are the prevalent method to approach the neural coding of sensory perception.^12,33,45^ Despite the success in accounting for the neural coding of lower-level acoustic-phonetic features,^12,13,37,46-48^ linear encoding models have not performed as well for higher-order speech information, and often fail to reveal information beyond acoustic stimulus encoding (Fig 5b, e). To account for the nonlinear transformations of the pure acoustic cues in the auditory system, higher-order features are included as predictors, such as phonetic, phonemic, syllabic, and lexical features.^13,37,44,49-51^ However, these feature representations rely on strong presumptions of hierarchical neural coding of these exact divisions and may not cover the intermediate representations in the non-primary auditory cortex.^52–54^ Furthermore, these models posit the auditory system as a passive finite response filter, which does not reflect the non-onset recurrent activity prevalent in high-order speech auditory areas.^37,52,55^

Traditional hierarchical models of neurobiology imply that specific brain areas are specialized for a particular representational level and that the transformation of information occurs across a anatomically defined “stream”, i.e. sound to phoneme to syllable to word and semantics.^11,56^ Our emerging findings challenge this traditional view. Instead, our results support that while there is a transformation from spectrogram to phonetic features, instead of phonemes and syllables as discretely encoded representations, we find complex, distributed higher-order representations that also carry forward prosodic information that may originate at earlier auditory levels and that processing is highly context dependent in later layers of computation. This may explain why we see not only phonetic feature tuning in STG,^13^ but also many “lower” level representations, such as onset, peakrate, frequency tuning,^37,50^ and “higher” level representations, such as context dependent, normalization, lexical effects.^15,44,49,51^

Our results revealed two critical factors determining the superior performances of the data-driven DNN models over the heuristic linear models with static speech feature sets. 1) The nonlinearity of the DNN models: specifically, almost all DNN layers consistently outperform the feature TRF models, even in the auditory periphery. This is indeed consistent with demonstrations of nonlinear processing in the auditory periphery.^57^ Given that both DNN and feature encoding models have roughly similar amounts of predictors (on the order of 100), this suggests that DNNs are learning nonlinear representations that are critical for extracting relevant phonetic features. 2) The dynamic temporal integration of contextual information in the DNN models: this is particularly critical for higher-order speech responses in the non-primary auditory cortex. Neural responses in STG were better predicted using deeper contextual representation layers in DNN with an extended delay time window. Furthermore, simply using nonlinear features in CNN layers with an even longer delay time window was not sufficient to achieve similar brain prediction performance for STG (Fig. 2). This indicates that specific dynamic temporal integration, which is aligned to the contextual information in speech and can be parametrized by computations such as Transformer-encoders or recurrent neural networks, is critical for characterizing neural responses to speech in STG. This finding provides evidence for STG speech processing units at dynamic timescales, which is possibly a mechanism for temporal binding of phonological sequences to form context-dependent acoustic-phonetic representations and ultimately perceptual representations of speech.^55^

Our results offer new perspectives for the underlying computations in the auditory pathway. A critical factor in deep neural network models is the model architecture, which determines the capability of computation and representation in the network.^58^ We found that different types of computational architectures were best correlated to different parts of the auditory pathway: the auditory periphery and subcortical areas were better characterized by locally-resolved static filters such as the convolution layers in DNN; the speech auditory cortex was better characterized by the sequential contextual representation layers, such as Transformer encoders and LSTM layers in DNN, which have more complex stimulus-dependent temporal dynamics than static spectrotemporal filters. These computations can be seen as signature features for different parts of the pathway: the auditory periphery and subcortical structures mainly consist of ascending feedforward synaptic connections, facilitating fast feedforward filtering of the signal;^35^ the speech auditory cortex, on the other hand, has multi-layer architecture with reciprocal connections that would facilitate sustained computations similar to recurrence and attentions.^59^ Of note, many previous studies have focused on cortex alone, suggesting that the main computations for speech along the path from primary to non-primary cortex. Here, through the lens of speech representation learning DNNs, we show that speech-relevant computations occur throughout the auditory pathway, and the periphery may play the more important computational role than primary auditory cortex.

This possibility has major implications for interpreting the functional roles of primary and non-primary auditory cortices. In particular, we found that dynamic computations that account for context-dependent representations did not contribute to predicting speech responses in the primary auditory cortex beyond the static convolutional filters (Figs. 2&4). On the contrary, the ability to predict the sustained neural response to speech in the non-primary auditory cortex STG is strongly correlated to dynamic computations in the neural networks (Figs. 2,3,4). This discrepancy is in line with a recent study that suggests phonological and complex sound processing in STG is fundamentally different from the tonotopic, narrow-tuned sound processing in primary auditory cortex.^37^ Indeed, the STG receives direct inputs from thalamic nuclei that are part of the non-tonotopic, non-lemniscal pathway,^60–62^ and does not appear to be dependent on the functional primary auditory cortex.^63^ The findings from this work suggest that representations found in the primary auditory cortex are not necessarily contributors to advanced computational models of speech processing, despite previous assumptions that it causally functions like the primary visual cortex in ventral stream object recognition processing.^56,64^

Our results also demonstrate that networks trained with self-supervised methods achieve equal or better performance in predicting speech responses in the cortex, compared to the more prevalent supervised models. The choice of training objective is critical for representation learning in deep neural networks. Previous works have found that supervised discriminant learning, such as word classification,^21,22^ leads to feature representation that correlates to auditory neural responses in the cortex. Our results are consistent with these findings. However, instead of using a discrete classification task, we show that a specific type of supervised sequence-to-sequence learning task, automatic speech recognition, can be used to provide the inductive bias towards neurally correlated feature representations for speech. Furthermore, self-supervised learning objectives, including contrastive and predictive learning, also yield similar representation and correspondence to the actual speech responses in STG. Supervised classification task has a direct link to categorical selectivity in higher-level visual areas, hence becoming a natural target for learning visual representations in the cortex.^24,25^ For naturalistic speech perception, on the other hand, previous studies do not support discrete selective coding for word forms in STG populations, but rather a collection of local populations tuned to multiple complex acoustic-phonetic cues and temporal landmarks in speech.^13,15,37,50,55,65-67^ Therefore, a single supervised task, such as word decoding, may not capture all the underlying computations and representations in STG. Self-supervised learning, on the other hand, is able to learn richer representations that are beyond the requirement of pure speech recognition, such as prosodic information, speaker identity etc. Our results found that fine-tuning using a supervised ASR task on top of the unsupervised training did not further improve the overall brain-encoding performance in STG. On the contrary, the brain-prediction performance for the non-primary auditory cortex decreased in the deep layers after supervised fine-tuning (Supplement Fig. 3).

From a computational modeling perspective, our results extend previous successes in the literature that employ deep neural networks as models of sensory systems.^24,68^ Several recent studies have taken a straight end-to-end approach and directly trained DNNs to predict neural responses.^69,70^ While this approach directly optimizes the brain-prediction performance, it may require a significant amount of data to train such deep models. For instance, the sequence-to-sequence DNN models we use here have around 90 million parameters and are trained with ~1000h of speech data to achieve competitive performance.^16,17,19^ It is not feasible to collect a similar amount of neural data within our clinical settings. Furthermore, due to the nature of intracranial recordings, we are only able to have a sparse sample, on the order of 100 electrodes, of the auditory cortex from each participant. As a result, the learned representations from a straight end-to-end optimization to brain activity may be biased by the individual difference in electrode sampling. Fundamentally, we take a transfer learning paradigm, pretraining DNNs without any neural data as inputs, and demonstrate that speech representations learned by these DNN models are also transferable to the neural coding in the human auditory pathway. It is worth pointing out that the deep neural networks used in this study are all trained on a completely independent dataset from the one used for neural signals. Moreover, the unsupervised models have not used any explicit information about the speech content or any linguistic knowledge of the speech. Unlike classical computational models of speech perception, such as TRACE,^11^ which assume a strict acoustic-phonetic-lexical hierarchy and explicit top-down inference, we find that acoustic-phonetic hierarchy also emerge from pure data-driven self-supervised models. The fact that such a self-supervised framework yields a similar and correlated representation hierarchy as the human auditory system suggests that the two systems may share similar computations that extract critical statistical structures in speech.

There are a few important limitations in our current approach. Our results suggest how different levels of speech representations emerge from hierarchical bottom-up recurrent or self-attentional operations and how these representations correlate to the auditory cortex. However, our models do not include top-down modules, nor do we include cortical coverage of areas beyond the auditory cortex, such as the frontal areas. Therefore, it remains to be delineated how other areas in the language network interact with the auditory cortex and if these interactions modulate local and populational representations of speech, and furthermore, to what extent can these interactions be characterized by our proposed framework. Aside from the scale of coverage, we have also limited ourselves to analyzing the temporal dynamics within individual electrodes. How feature representations in deep neural networks are aligned to the distributed population-level neurodynamics^71^ in the auditory cortex remains to be investigated in future work.

A potential limitation is the biological plausibility of the computational models used in this study. We focused on the learned feature representations rather than the actual parametrization and implementations of the computing algorithm such as self-attention or long short-term memory mechanism. It is of course hard to claim that any of these computations are actually implemented in the cortex, or the gradient-based learning rule is adopted by the brain. However, it is promising is that the in silico models converge on a similar representational basis of speech sequences as the auditory cortex, with a learning algorithm that does not require millions of labeled examples and has been shown to be a potentially strong candidate for a biologically plausible theory of sensory learning,^29^ or higher-level language processing in general.^30^

In sum, using a comparative approach, we show important representational and computational similarities between speech learning DNNs and the human auditory pathway. From a neuroscientific perspective, our results demonstrate that data-driven computational models can extract relevant intermediate features from statistical structure of speech that outperform traditional feature-based encoding models. Furthermore, our results challenge the concept of auditory ventral stream which is based on hierarchical feedforward processing across successive cortical areas, but suggests contextual and time dependent processes are more important. On the other hand, from the deep neural network perspective, we provide a new avenue to open up the “black box” representations of DNNs by comparing them with neural responses and selectivity. We show that modern DNNs may have converged on algorithms and representations that approximate processing in the human auditory system.

## Methods

The experimental protocol was approved by the Institutional Review Board at the University of California, San Francisco (UCSF) and by the Huashan Hospital Institutional Review Board of Fudan University. All participants gave their written, informed consent prior to testing. All patient data were stored and analyzed on computing servers within UCSF, and Facebook AI Research only performed DNN model pre-training using publicly available speech corpora, without any access to the patient data.

### Participants

This study included 12 monolingual participants (6 Male, 6 Female, age from 31 to 55, all right-handed) who were neurosurgical patients at either UCSF Medical Center or Huashan Hospital. The nine native English-speaking participants from UCSF (E1-E9) were either eloquent brain tumor patients (4 patients) undergoing awake language mapping as part of their surgery or patients with intractable epilepsy (5 patients) who had high-density electrode grids implanted for clinical monitoring of seizure activity (all left hemisphere coverage). We only included those participants with tumors that did not obviously invade the auditory cortex. Three native Mandarin-speaking participants from Huashan Hospital (M1-M3) were eloquent brain tumor patients undergoing awake language mapping as part of their surgery (3 left hemisphere coverage). The placements of the grids were determined solely by clinical needs. All patients were clearly informed (as detailed in the IRB-approved written consent document signed by the participant) that the participation in the scientific research was completely voluntary and would not directly impact their clinical care. Additional verbal consent was also acquired at the beginning and during the breaks of each experiment session.

### Experiment paradigm

During the experiments, the participants were instructed to passively listen to continuous speech stimuli. The acoustic stimuli used in this study consisted of natural, continuous speech in both American English and Mandarin. The English speech stimuli consisted of materials from the TIMIT dataset.^39^ The TIMIT set consisted of 499 English sentences selected from the TIMIT corpus, spoken by 402 different speakers (286 males and 116 females). The sentences were separated with 0.4 sec of silence. The task was broken into 5 blocks with each block ~5 min in time. The Mandarin speech was a subset of the Annotated Speech Corpus of Chinese Discourse (ASCCD) from the Chinese Linguistic Data Consortium,^72^ which included read texts of a variety of discourse structures, such as narrative and prose. The stimuli set consisted of 68 passages of Mandarin speech selected from the ASCCD corpus, spoken by 10 different speakers (5 males, 5 females). The length of single passage varied between 10 to 60 sec. The passages were separated with 0.5 sec of silence. The task was broken into 6 blocks with each block ~5 min in time.

Depending on their clinical conditions, each participant finished 3 to 11 blocks of all tasks. In particular, 8 English speaking participants (E1-E8) finished all 5 TIMIT blocks, E9 finished 3 TIMIT blocks, and 3 Mandarin speaking participants (M1-M3) finished 2 TIMIT blocks. 3 English speaking participants (E1-E3) and 3 Mandarin speaking participants (M1-M3) finished all 6 ASCCD blocks. E4 finished 5 ASCCD blocks.

### Data acquisition and preprocessing

In all patients, the same types of high-density ECoG grids (manufactured by Integra or PMT) with identical specifications (4 mm center-to-center spacing and 1.17 mm diameter exposed contact lateral) were placed on the lateral surface of the temporal lobe. Depending on the exact clinical need, the grid may have 32 (8 x 4), 128 (16 x 8) or 256 (16 x 16) contact channels in total. In 4 patients (E6-E9), an extra 32 channel (8 x 4) grid with 4mm center-to-center spacing and 1.17 mm diameter exposed contact lateral grids (Integra) was placed on the temporal plane for each patient. During experimental tasks, neural signals were recorded from the ECoG grids using a multichannel amplifier optically connected to a digital signal processor (Tucker-Davis Technologies). The TDT OpenEx software was used for data recording. The local field potential at each electrode contact was amplified and sampled at 3052Hz. The raw voltage waveform was visually examined, and channels containing signal variation too low to be detectable from noise or continuous epileptiform activity were removed. Time segments on remaining channels that contained electrical or movement-related artifacts were manually marked and excluded. The signal was then notch-filtered to remove line noise (at 60Hz, 120Hz, and 180Hz for English-speaking participants and at 50Hz, 100Hz, and 150Hz for Mandarin-speaking participants) and re-referenced to the common average across channels sharing the same connector to the preamplifier.

Using the Hilbert transform, the analytic amplitude of eight Gaussian filters (center frequencies: 70-150Hz) was computed. The high-gamma signal was taken as the average analytic amplitude across these eight bands. The signal was down-sampled to 100Hz. The tasks were broken into recording blocks of ~5 minutes in length. The high-gamma signal was z-scored across the recording block.

### Electrode localization

For the chronic monitoring cases, electrodes were localized by aligning preimplantation MRI and post-implantation CT scans. For the awake cases, high-density electrode grids were temporarily placed onto the temporal lobe intraoperatively to record cortical local potentials. The three-dimensional positions of the corners of the grid were recorded using the Medtronic neuronavigation system and then aligned to the pre-surgery MRI. Intraoperative photographs were used as references. The remaining electrodes were localized using interpolation and extrapolation from those points.^73^

### Data analysis software

All analyses were carried out using custom software written in Python and Matlab. Custom Matlab code was used for data preprocessing. Open-source scientific Python packages used included pytorch, fairseq, huggingface transformers, numpy, scipy, pandas, librosa, and scikit-learn. Cortical surface reconstruction was performed using Freesurfer and electrodes were co-registrated using Python package img-pipe. Praat^74^ was used to extract pitch features. Figures were created with matplotlib and seaborn in Python.

### Biophysical models for auditory peripheral and mid-brain

We used neuronal models of the midbrain and auditory periphery.^34–36^ The model consists of a phenomenological model of the auditory nerve (AN) responses, which includes nonlinear properties such as rate saturation, adaptation, and synchrony capture and an extended same-frequency inhibition-excitation model of inferior colliculus (IC), which includes both band-pass and low-pass/band-reject IC cells. The synaptic outputs from 50 auditory nerve neurons with characteristic frequency uniformly distributed on log scale within [150, 8000] hz were extracted as the AN signal. These synaptic output signals were used as inputs to the two different types of midbrain neurons in area IC, which resulted in 50 band-pass IC neurons and 50 low-pass/band-reject IC cells.

For each speech sentence, the raw waveform was sent into the model as input, and the corresponding response sequences from AN and IC cells were extracted and down-sampled to 100 hz to match the high-gamma signals from the cortex.

### Acoustic, phonetic and prosodic feature definitions

We use a heuristic set of 208 features as the baseline prediction model (161 spectrum, 13 phonetics, 31 pitch/prosodic, 3 envelope).

The spectrogram features of the speech were calculated using short-time Fourier transform, with 161 frequency components ranging from 0 to 8KHz in log-scale.

The phonetic features were 13-dimensional binary time series similar to previous works.^13,37^ These features describe single phonemes as a combination of the places of articulation (dorsal, coronal, labial), manners of articulation (plosive, fricative, nasal), and voicing of consonants, as well as vowel place (high, mid, low, front, back), and indicator of consonant/vowel.

The pitch features were extracted in the same way as in our previous work,^44^ including absolute pitch, speaker-normalized relative pitch and pitch change, and a binary variable when pitch values were present, indicating voicing in the speech. The fundamental frequency (F0) was calculated using an automated autocorrelation method in Praat and corrected for halving and doubling errors. The absolute pitch was defined as the natural logarithm of F0 values in Hz. The relative pitch was computed by z-scoring the absolute pitch values (log F0) within each sentence/passage (within speaker). The pitch change was computed by taking the first-order derivative (finite difference) in time for logF0. We discretized absolute pitch, relative pitch and pitch change into 10 bins, equally spaced from the 2.5 percentile to the 97.5 percentile value. The bottom and top 2.5% of the values were placed into the bottom and top bins respectively. As a result, absolute pitch, relative pitch and pitch change were represented as three 10-dimensional binary feature vectors. For non-pitch periods, these feature vectors would have all 0s.

The envelope features included intensity, sentence onsets, and peak rates. Intensity is a continuous scalar sequence representing the envelope of the speech. The sentence onset is a binary feature with a 1 at the onset of the first time stamp of the first phoneme in each sentence, and 0 elsewhere. The peak rate was computed in the same way as previous work,^50^ which was a sparse time series of local peaks extracted in the first order derivative of the amplitude envelope of the speech.

### Encoding models

We used time-delayed linear encoding models known as temporal receptive field models.^12^ Temporal receptive field (TRF) models allow us to predict neural activity based on stimulus features in a window of time preceding neural activity. In particular, we fit the linear model 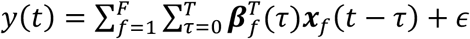 for each electrode, where *y* is the high-gamma activity recorded from the electrode, ***x**_f_*(*t* – *τ*) is the stimulus representation vector of feature set *f* at time *t- τ*, ***β**_f_*(*τ*) is the regression weights for feature set *f* at time lag *τ*, and *ϵ* is the gaussian noise.

To prevent model overfitting, we used L2 regularization and cross-validation. Specifically, we divided the data into three mutually exclusive sets of 80%, 10% and 10% of samples. The first set of 80% was used as the training set. The second set was used to optimize the L2 regularization hyperparameter, and the final set was used as the test set. We evaluated the models using the correlation between actual and predicted values of neural activity on held out data. We performed this procedure 5 times and the performance of the model was taken as the mean of performance across all testing sets.

The performance of each specific encoding model on an individual recording site (electrode/neuron) was quantified as the (normalized) brain prediction score (BPS). In particular 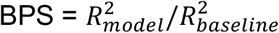, where 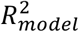 is the *R*^2^ value of the prediction model based on cross-validation, and 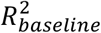 is the *R*^2^ value of the baseline model (full feature set model) for the same electrode/neuron based on cross-validation. A BPS value of 1 indicates the proposed model performs as good as the baseline, and BPS larger than 1 suggests the proposed model outperforms the baseline model.

For the spectrogram TRF (STRF) model and the baseline full feature model, we used a fixed 400 ms delay time window length. For all the DNN-based encoding models, we varied the time window length from 0 (only using the current time frame) to 400 ms, and picked the optimal window length based on the cross-validation results.

### Electrode selection

To select speech responsive electrodes, and to avoid numerical instability of BPS caused by dividing very small *R*^2^ values of baseline, we only include speech-responsive electrodes in our analysis. The responsive threshold was set as 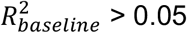.

### Deep neural networks: model architectures

We used 4 different DNN models, namely HuBERT (HuB),^19^ Wav2Vec 2 unsupervised version (W2V),^17^ Wav2Vec 2 ASR-supervised (W2V-A),^17^ and DeepSpeech 2 (DS2).^16^

The HuBERT and Wav2Vec 2 models shared the same architecture, consisting of a convolutional waveform encoder and a Transformer BERT encoder.^75^ The network took 16kHz raw sound waveform as input. The convolutional encoder consisted of seven 512-channel 1D convolution layers with strides [5, 2, 2, 2, 2, 2, 2] and kernel widths [10, 3, 3, 3, 3, 2, 2]. The convolution encoder down-sampled the input to 512-dimensional feature sequence at 20ms framerate (50Hz). The output of the convolution encoder, noted as “CNN out”, was projected to 768 dimensional space through a linear layer, noted as “CNN proj”, and fed into the BERT encoder. The BERT encoder consisted of 12 identical transformer encoder blocks, with embedding dimension of 768, intermediate feedforward layer dimension as 3072, and 12 attention heads in each layer.

The DeepSpeech 2 model consisted of a convolutional spectrogram encoder and a recurrent encoder. The DS2 model took in the spectrogram of the raw audio signal as input. The spectrogram was computed using a short-time Fourier transform with 161 frequency components from 0 to 8kHz, time window size of 0.02s and a stride size of 0.01s. The convolution encoder consisted of two 32-channel 2D convolution layers, with 2D strides [2, 2] and [2, 1] correspondingly, and kernel size (41, 11) and (21, 11) correspondingly. The final output of the convolution encoder was a 1312-dimensional vector at 20ms framerate (50Hz). The recurrent encoder consisted of 5 bi-directional long short-term memory (LSTM) layers, each with hidden state size of 1024. The output of the last LSTM layer was projected to 29-dimensional feature space by a linear projection layer.

### Deep neural networks: unsupervised training

The HuBERT model was trained using a self-supervised paradigm of masked prediction.^19^ The unsupervised k-means clustering algorithm was used to generate categorical labels of the acoustic speech signal, mimicking the pseudo-phonetic labels. During training, a random subset of segments in each sentence were selected and masked. After masking, the sequence was passed through the network to generate a feature embedding sequence. The embedded sequence was then projected to compute cross-entropy loss over the discrete code categories.

The Wav2Vec 2 unsupervised model was trained using a self-supervised contrastive learning paradigm.^17^ The model used a quantization module to discretize the output sequence of the convolution encoder. Similar to BERT, a random subset of speech segments were selected and masked. The final output of the Transformer encoder and the quantized representation from the convolution encoder were used to compute the contrastive loss. Specifically, for the target output at a given masked time step, a random set of distractors were picked from other masked portions in the same sentence. The contrastive loss maximizes the distance between the target and the discretized output in the distractors, while minimizing the distance between the target and the discretized output at the target time step.

Both models were trained on the 960h Librispeech corpus.^40^ For the cross-language comparison, we also trained a HuBERT Mandarin model on 755h MAGICDATA corpus of Mandarin speech^76^ using the same procedure.

### Deep neural networks: supervised training

The Wav2Vec 2 supervised model was fine-tuned from the unsupervised pre-trained initialization.^17^ A linear projection layer was used to project the output of the Transformer encoder onto 29 classes representing characters, space, and word boundary. The model was optimized by minimizing a connectionist temporal classification (CTC) loss.^77^ During fine-tuning, the weights of the convolution encoder were frozen.

The DeepSpeech 2 model was trained from random initialization for best automatic speech recognition (ASR) performance by minimizing the CTC loss.^16^ The supervised training of both models used the 960h LibriSpeech corpus.

### Attention pattern analysis

For a given speech sentence, assume the embedding sequence in a Transformer layer was of length T [*c_1_*,…, *c_T_*], and phoneme boundaries indexed as [*p_1_*,…, *p_m_*], syllable boundaries indexed as [*s_1_*,…, *s_n_*]. The attention templates were defined as follows:

1. Attention to the current phoneme, phoneme (0): 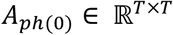, *A*_*ph*(0)_(*i,j*) = 1 if *p_k_* ≤ *i* < *P*_*k*+1_ and *P_k_* ≤ *j* < *P*_*k*+1_ for any *k, A*_*ph*(0)_(*i,j*) = 0 otherwise.
2. Attention to the previous phoneme, phoneme (−1): 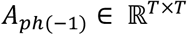, *A*_*ph*(−1)_(*i,j*) = 1 if *P_k_* ≤ *i* < *P*_*k*+1_ and *P*_*k*-1_ ≤ *j* < *P_k_* for any *k*, *A*_*ph*(−1)_(*i,j*) = 0 otherwise.
3. Attention to the second to previous phoneme, phoneme (−2): 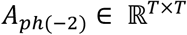, *A*_*ph*(−2)_(*i,j*) = 1 if *P_k_* ≤ *i* < *P*_*k*+1_ and *P*_*k*-2_ ≤ *j* < *P*_*k*-1_ for any *k*, *A*_*ph*(−2)_(*i,j*) = 0 otherwise.
4. Attention to the current syllable, syllable (0): 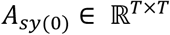, *A*_*sy*(0)_(*i,j*) = 1 if *s_k_* ≤ *i* < *s*_*k*+1_ and *s_k_* ≤ *j* < *s*_*k*+1_ for any *s*, *A*_*ph*(0)_(*i,j*) = 0 otherwise. And to exclude the current phoneme from the current syllable, we use 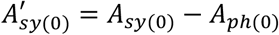 as the template.
5. Attention to the previous syllable, syllable (−1): 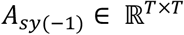, *A*_*sy*(−1)_(*i,j*) = 1 if *s_k_* ≤ *i* < *s*_*k*+1_ and *s*_*k*-1_ ≤ *j* < *s_k_* for any *k*, *A*_*sy*(−1)_(*i,j*) = 0 otherwise.
6. Attention to the second to previous syllable, syllable (−2): 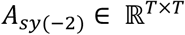, *A*_*sy*(−2)_(*i,j*) = 1 if *s_k_* ≤ *i* < *s*_*k*+1_ and *s*_*k*-2_ ≤ *j* < *s*_*k*-1_ for any *k*, *A*_*sy*(−2)_(*i,j*) = 0 otherwise.

For the sentence, we computed the attention matrix *W_xy_* at the *x*-th layer and *y*-th attention head. The correlation coefficient *corr*(*W_xy_, A_q_*) was computed for all templates. And attention score (AS) for layer *x* and template *q* was computed as the average over all attention heads and over all speech sentences.

### STG clustering analysis

To identify functional clusters in STG, we took a similar clustering approach as described in previous works.^41^ Specifically, we applied convex non-negative matrix factorization (convex NMF)^78^ to decompose the averaged high-gamma time series across all STG electrodes. Specifically, 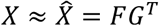, and *F* = *XW*, where *X* [*T* time points × *p* electrodes] was the ERP matrix for different STG electrodes average across all sentences, *G* [*p* electrodes × *k* clusters] represented spatial weight of each electrode for each cluster, and *W* [*p* electrodes × *k* clusters] represents weights applied to electrode time series. In particular, for *X* we took all 144 speech-responsive STG electrodes across all 9 subjects and computed the averaged ERP response for each electrode across all 599 TIMIT sentences. We evaluated different *k* values ranging from 1 to 10 and computed the percent of variance explained by NMF models with different *k* values. We chose the number of clusters at the elbow of the percent variance curve (Supplement Figure 4), which yielded *k* = 2, and explained 94% of the total variance.

After choosing the optimal number of clusters, each electrode was assigned to a cluster with the maximum cluster weight in *G*.

## Acknowledgments

We would like to thank Matthew Leonard, Laura Gwilliams, Ilina Bhaya-Grossman, Yizhen Zhang and Laurel Carney for critically reading the manuscript and for helpful suggestions. The authors gratefully acknowledge the support of the National Institute of Neurological Disorders and Stroke under U01NS117765 (to E.F.C.), the National Institute on Deafness and Other Communication Disorders under R01DC012379 (to E.F.C.), the William K. Bowes Foundation (to E.F.C.), the William and Susan Oberndorf Foundation (to E.F.C), the Joan and Sanford Weill Foundation (to E.F.C.), the Shurl and Kay Curci Foundation (to E.F.C.), Shanghai Municipal Science and Technology Major Project under No.2018SHZDZX01 (to J.W.), Shanghai Shenkang Hospital Development Center under SHDC12018114 (to J.W.), Shanghai Rising-Star Program under No. 19QA1401700 (to J.L.) and Shanghai Young Talents Program under No. 2017YQ014 (to J.L.).

## Author Contributions

Conceptualization: Y.L. and E.F.C.; Methodology: Y.L., G.K.A., and A.M.; Software: Y.L., G.K.A., and A.M.; Formal analysis: Y.L.; Resources: E.F.C., J.W. and J.L.; Writing – Original Draft: Y.L.; Writing – Review & Editing: Y.L., G.K.A., A.M. and E.F.C.; Supervision: E.F.C.

---

**Supplement Figure 1.**
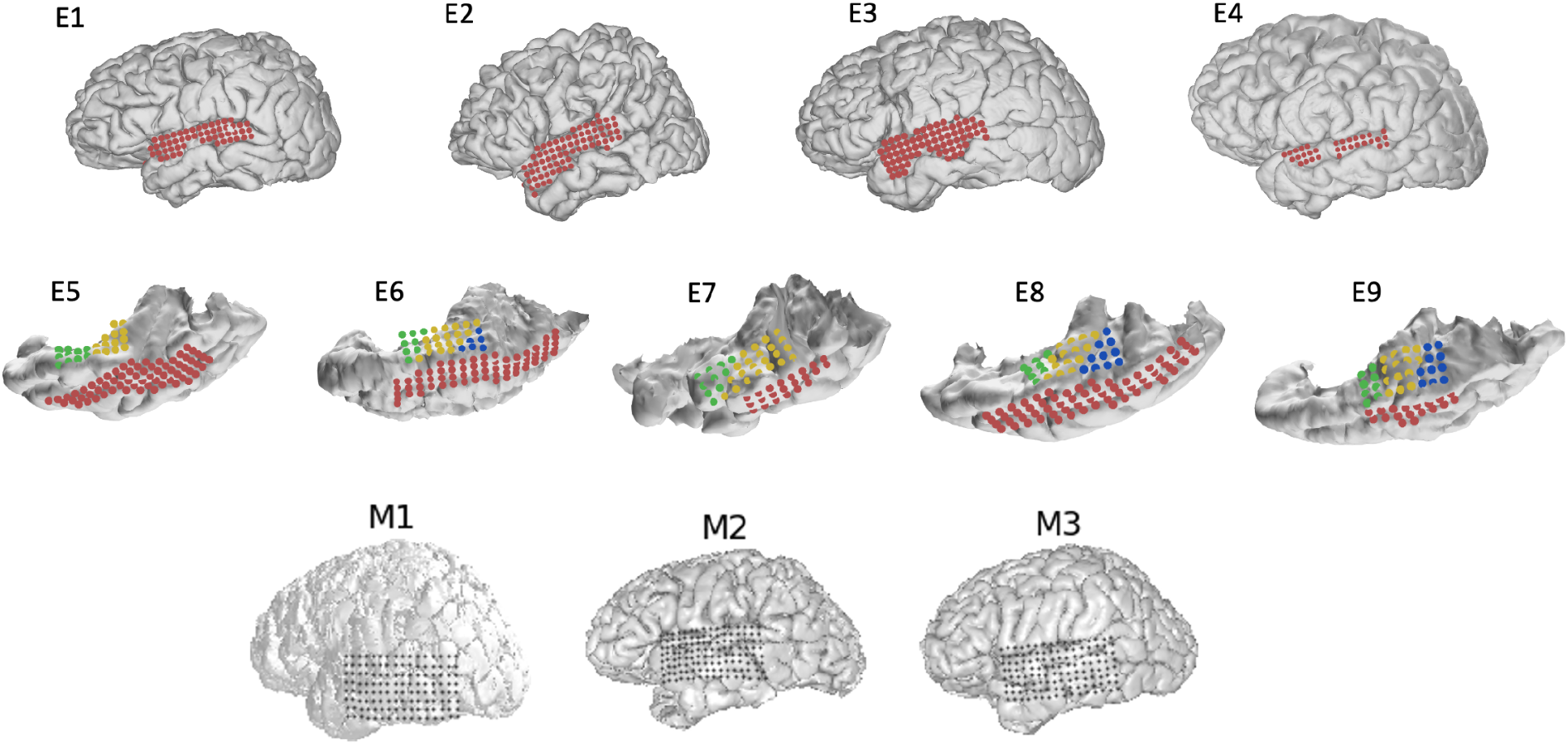
ECoG grid coverage for all subjects. For E1-E9, electrodes are marked in colors according to anatomical label: superior temporal gyrus (red), Heschl’s gyrus (yellow), planum temporale (blue), planum polare (green).

**Supplement Figure 2.**
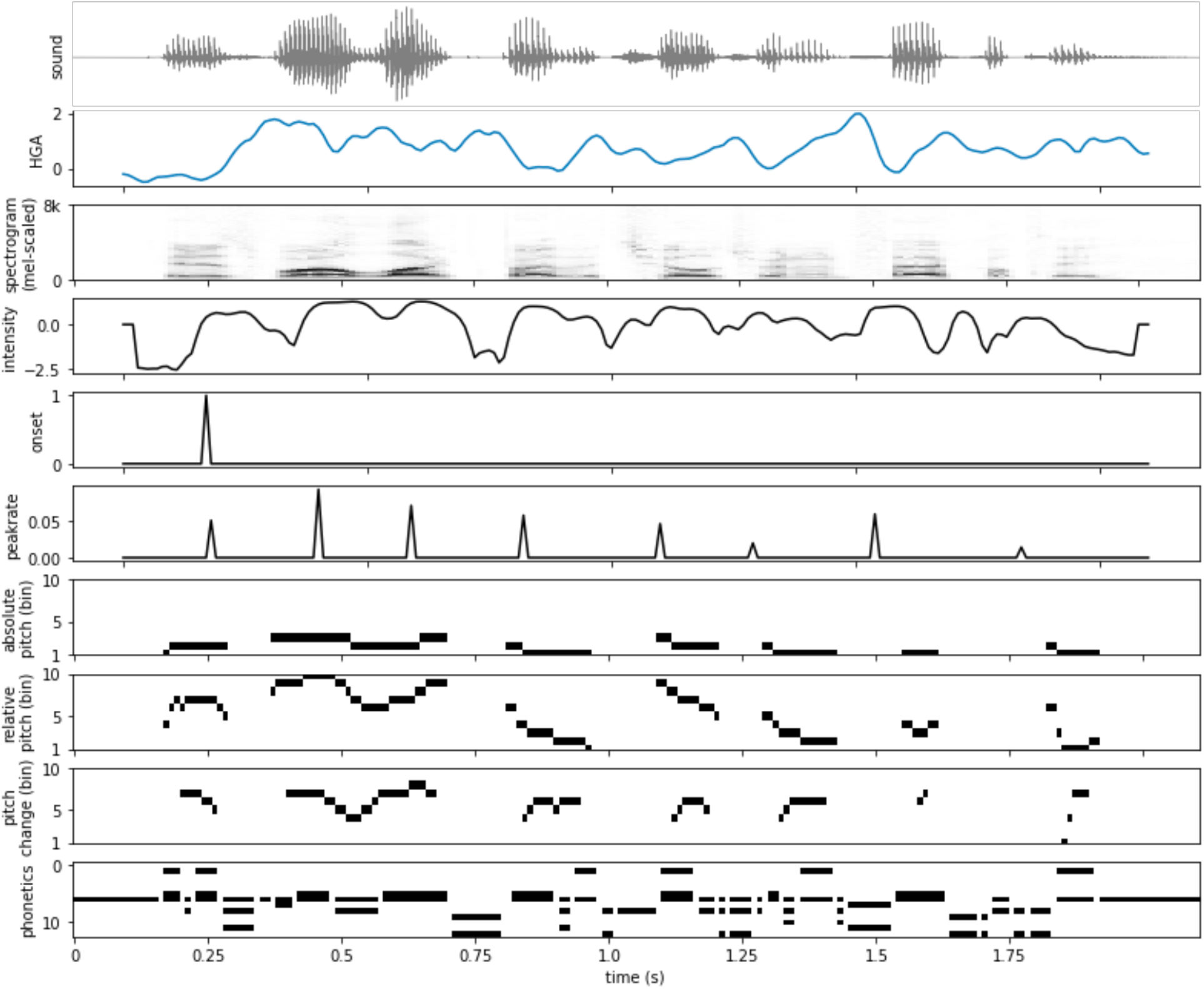
Acoustic phonetic feature encoding model. Example of feature extraction for a sample sentence. From top to bottom: 1) raw waveform; 2) high-gamma (z-scored) activity at an example electrode; 3) Mel-scaled spectrogram; 4) intensity of voicing; 5) sentence onset; 6) time course of peak rate; 7) absolute pitch (binned into 10 bins); 8) relative pitch (binned into 10 bins); 9) pitch change (binned into 10 bins); 10) phonetic features.

**Supplement Figure 3.**
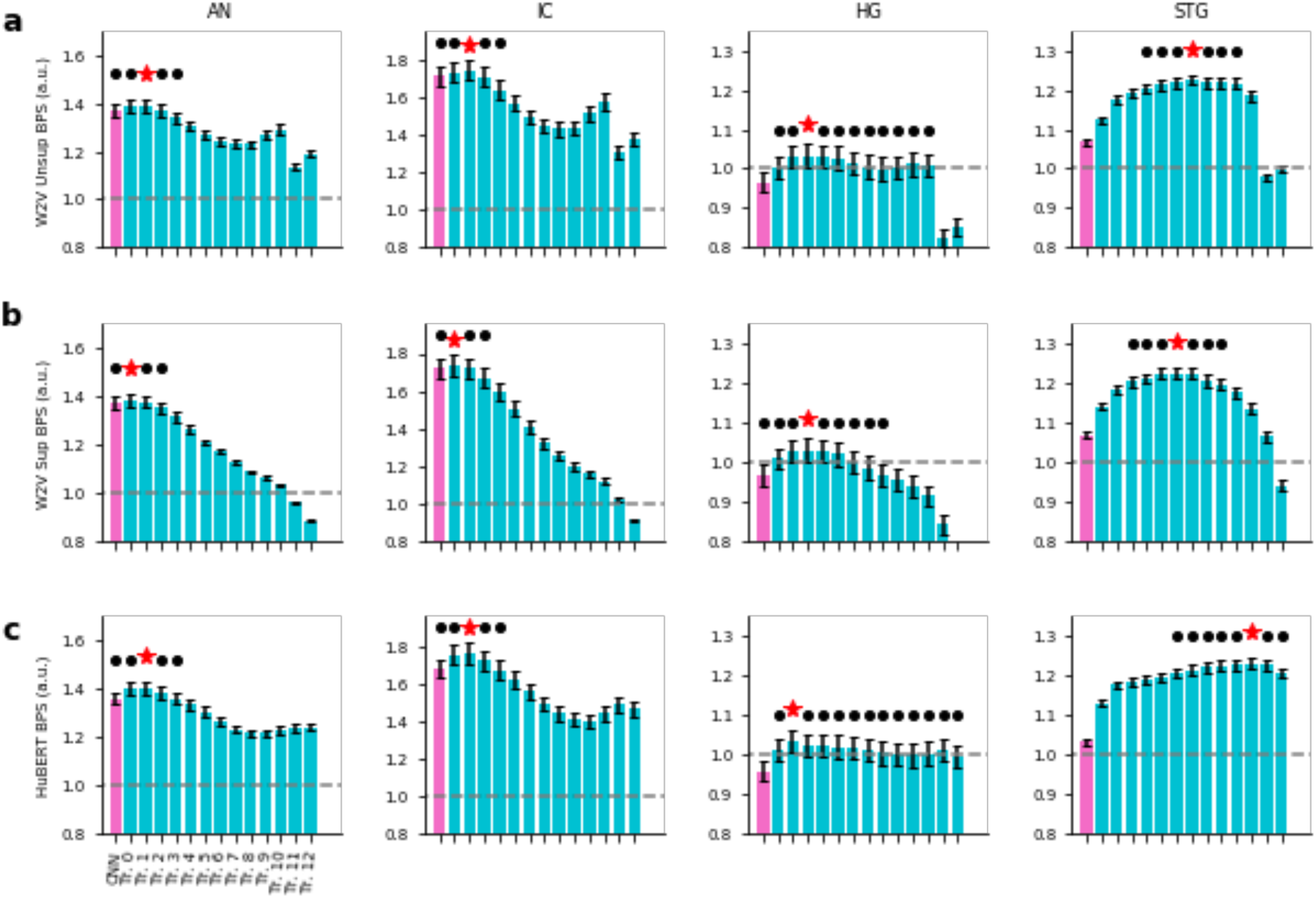
Comparing DNN encoding performance across different models. The averaged normalized brain prediction score of the best-performing neural encoding model based on each single layer in the DNN model (maximum over delay window length). **a**) Wav2Vec 2.0 Unsupervised model; **b**) Wav2Vec 2.0 Supervised model; **c**) HuBERT model. Each column corresponds to one area in the auditory pathway, from left to right AN/IC/HG/STG. Magenta bars indicate CNN output layers, cyan bars indicate Transformer layers. Red star (*) indicates the best model for each area, black dot (.) indicates other models that are not statistically different from the best model (p > 0.05, two-sided paired t-test).

**Supplement Figure 4.**
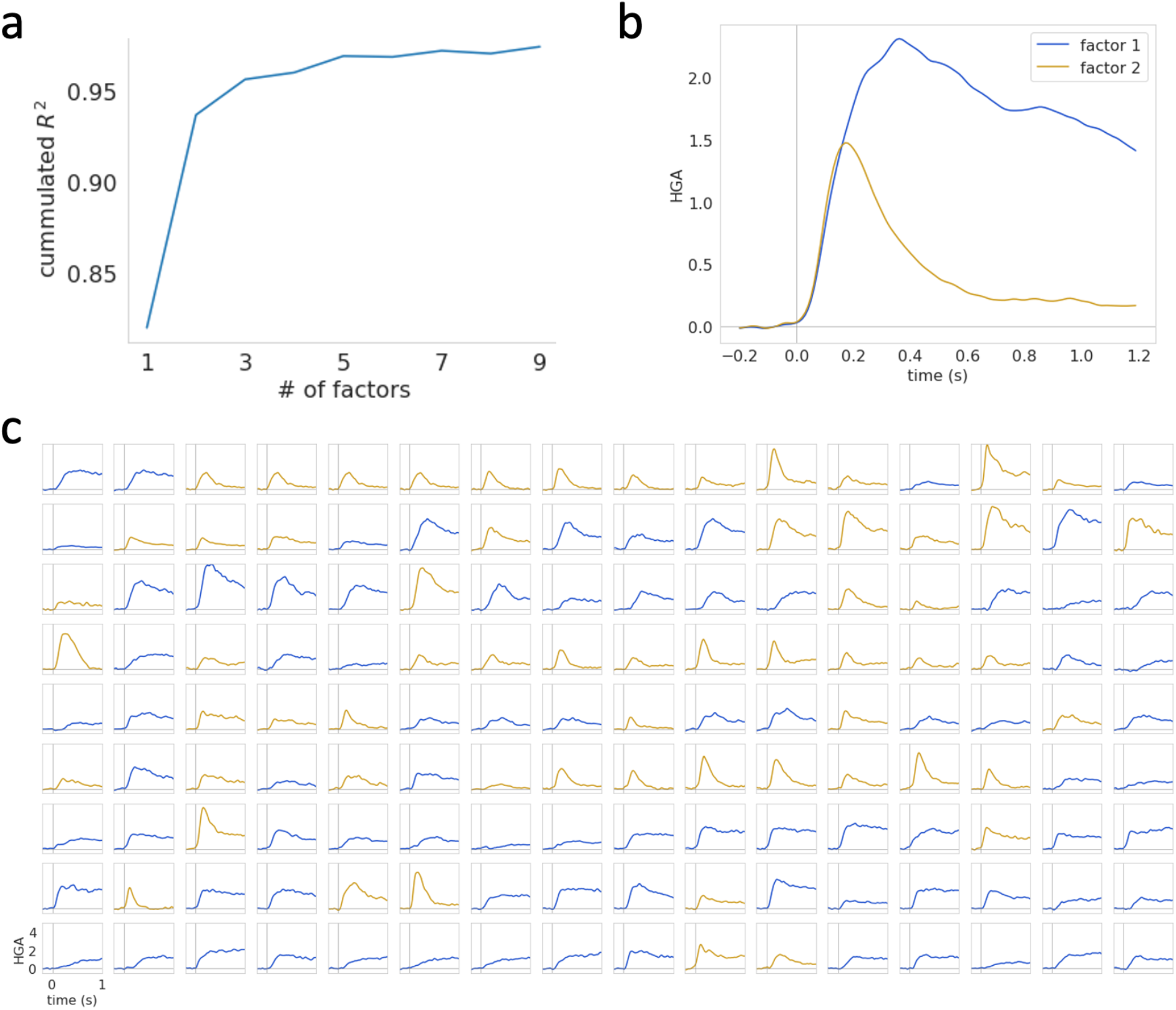
Clustering the STG electrodes. **a**) Percent of total variance explained by the NMF decomposition with different number of factors; **b**) The time course of the two factors from the NMF model; **c**) the cluster assignment for each STG electrode. Each panel is the sentence averaged response for one STG electrode.

